# Metabolites that feed upper glycolytic branches support glucose independent human cytomegalovirus replication

**DOI:** 10.1101/2024.02.12.579992

**Authors:** Rebekah L. Mokry, John G. Purdy

## Abstract

The broad tissue distribution and cell tropism of human cytomegalovirus indicates that the virus successfully replicates in tissues with various nutrient environments. HCMV requires and reprograms central carbon metabolism for viral replication. However, many studies focus on reprogramming of metabolism in high nutrient conditions that do not recapitulate physiological nutrient environments in the body. In this study, we investigate how HCMV successfully replicates when nutrients are suboptimal. We limited glucose following HCMV infection to determine how glucose supports virus replication and how nutrients potentially present in the physiological environment contribute to successful glucose independent HCMV replication. Glucose is required for HCMV viral genome synthesis, viral protein production and glycosylation, and virus production. However, supplement of glucose-free cultures with uridine, ribose, or UDP-GlcNAc—metabolites that support upper glycolytic branches—resulted in partially restored viral genome synthesis and subsequent partial restoration of viral protein levels. Low levels of virus production were also restored. Supplementing lower glycolysis in glucose-free cultures using pyruvate had no effect on virus replication. These results indicate nutrients that support upper glycolytic branches like the pentose phosphate pathway and hexosamine pathway can compensate for glucose during HCMV replication to support low levels of virus production. More broadly, our findings suggest that HCMV could successfully replicate in diverse metabolic niches, including those in the body with low levels of glucose, through alternative nutrient usage.

**IMPORTANCE:** The metabolic environment is a determinant in the ability of a virus to successfully replicate. HCMV has broad cell tropism and replicates in various tissues that have diverse and/or limiting metabolic environments. We know that HCMV reprograms host central carbon metabolism to support viral replication, but we have little understanding of HCMV replication in diverse metabolic niches as most studies use high nutrient culture media. Here, we show that glucose limitation suppresses virus production through loss of viral genome synthesis and viral protein glycosylation. However, nutrient compensation by uridine, ribose, and UDP-GlcNAc, metabolites that fuel upper glycolytic branches such as the non-oxidative pentose phosphate pathway support low levels of glucose-independent virus production. Our work indicates that metabolite compensation may facilitate HCMV replication in nutrient limited niches in the body.

## INTRODUCTION

Human cytomegalovirus (HCMV) is a prevalent herpesvirus that causes severe disease in immunocompromised individuals (1–3). The virus can cause a range of diseases in affected individuals, and common complications include CMV pneumonitis, retinitis, and hepatitis (2–4). Initiation of CMV diseases are facilitated through the virus’s ability to infect, replicate, and spread in broad and diverse tissue via its extensive cell tropism (2). HCMV requires host metabolism for replication and promotes and reprograms central carbon flow to obtain components required for viral replication, such as nucleotides for viral genome replication, amino acids for viral protein production, UDP-sugars for viral protein glycosylation, and lipids for the viral membrane (5–18).

Glucose is an important carbon supplier for viral replication components and is essential for optimal infectious virus production in culture (19, 20). HCMV promotes glucose uptake (7, 21, 22), which is accompanied by increased glycolytic flux during infection (8–10). Glucose supports key branch points in upper glycolysis such as the pentose phosphate pathway (PPP) and ribose 5-phosphate production via glucose 6-phosphate, and the hexosamine pathway and UDP-N-acetylglucosamine (UDP-GlcNAc) production via fructose 6-phosphate (9, 10, 23, 24). Glucose carbons are also shuttled to citrate and then to malonyl-CoA for fatty acid synthesis and elongation and lipid synthesis (9, 10, 17, 18, 25, 26). Chemical inhibition of glucose uptake, glycolysis, or the PPP (21, 23, 24, 27) as well as suppression of glycolytic activating proteins (22, 24, 28) result in loss of infectious virus production.

To date, investigations examining glucose carbon flow in HCMV infection have used well defined cell culture media conditions. However, one caveat to these metabolic investigations in culture is that some nutrients are at supraphysiological levels while others are absent (29, 30). Use of these media could mask the contribution of alternative metabolites to HCMV replication, particularly those that provide essential contributions in nutrient poor conditions. Currently, it is unknown how HCMV successfully replicates in nutrient poor environments that lack or limit major metabolites required for viral replication. Here, we investigate the impact of glucose deprivation on HCMV replication and alternative metabolite compensation that promotes HCMV replication in glucose limited environments. We demonstrate that glucose deprivation attenuates viral genome synthesis, and subsequently, viral protein production and viral protein glycosylation.

Metabolite supplement using uridine or ribose, which feed upper glycolytic branches, partially restore viral genome synthesis. Importantly, these upper glycolytic metabolites enable low levels of virus production independent of glucose. Additionally, ribose also rescued viral protein glycosylation at late replication stages, suggesting it contributes to the production of multiple components for viral replication. Further, UDP-GlcNAc, a product of the upper glycolytic branch hexosamine pathway and the substrate for both O- and N-linked glycosylation, provides a subtle increase in viral protein glycosylation while partially restoring viral genome levels and low levels of virus production.

Supplementing with pyruvate, the final product of glycolysis, did not restore viral genome levels or virus production. Overall, these studies reveal that metabolic compensation from upper glycolytic branches allows low yet persistent virus production in glucose limiting environments.

## RESULTS

### Glucose deprivation attenuates HCMV genome synthesis, resulting in decreased viral protein levels and inhibition of infectious virus production

Glucose deprivation inhibits infectious virus production in HCMV-infected cells (19, 20). However, it is unknown which stage of replication is impacted by the loss of glucose. We examined multiple steps of HCMV replication during glucose deprivation to determine which stages require glucose. Human foreskin fibroblasts (HFF) were infected with HCMV strain TB40/E encoding green fluorescent protein (TB40/E-GFP). Experiments were performed in growth arrested, serum starved conditions to remove any contribution of nutrients from serum. These conditions are commonly used in metabolic studies of HCMV (8–10, 17, 18, 20–26, 31–33). Glucose was removed at 1 hour post infection (hpi) and virus production, viral genome synthesis, and viral protein levels were quantified (**Fig. 1**). Cell-free and cell-associated viral titers were measured using 50% tissue culture infectious dose assay (TCID_50_). Glucose-replete (25 mM) DMEM cultures had measurable levels of infectious cell-free virus production from 96-120 hpi and cell-associated virus production from 72-120 hpi with the highest titers at 120 hpi (**Fig. 1A**). Virus production was not detected in glucose-free DMEM cultures, consistent with previous studies (19, 20). To determine if loss of glucose decreases HFFs viability, we measured cytotoxicity of glucose-free conditions. Cells were infected or mock-infected as described above. At 48 and 120 hpi, culture supernatants were collected for lactate dehydrogenase cytotoxicity assay. At 48 hpi, cells maintained high viability in both mock and HCMV-infected cultures (**Fig. 1B**). By 120 hpi, mock-infected, glucose-free cell viability had decreased to 78%, while HCMV-infected, glucose-free cultures maintained 95% cell viability. The viability of HCMV-infected, glucose-replete cultures decreased to 89%, likely due to cell death that occurs from high levels of virus production at late stages of HCMV replication. These data are consistent with prior studies in HFFs (19) and indicate that HCMV offers protection against cell death induced by glucose deprivation.

**Fig 1.**
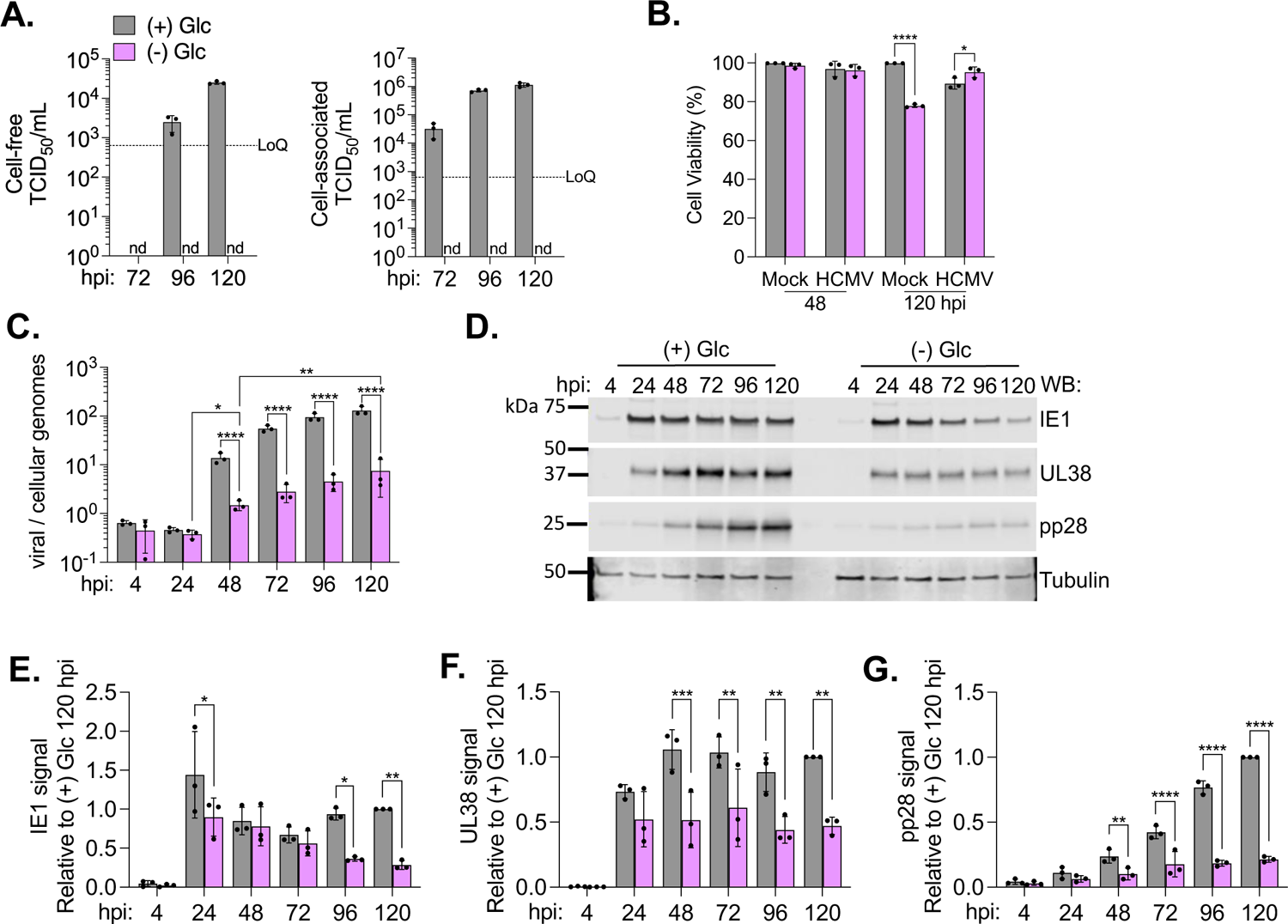
Glucose deprivation attenuates HCMV genome synthesis, viral protein levels, and inhibits virus production. Human foreskin fibroblasts (HFFs) were infected with HCMV strain TB40/E encoding free green fluorescent protein (TB40/E-GFP) at a multiplicity of infection (MOI) of 2 infectious units per cell (IU/cell). At 1 hour post infection (hpi), media was changed to glucose-replete [(+) Glc] or glucose-free [(-) Glc] DMEM. (**A**) At the indicated times, culture supernatant or cells were collected for cell-free (left) and cell-associated (right) virus production. Viral titers were measured by 50% tissue culture infectious dose assay (TCID_50_). LoQ = 632 TCID_50_/mL; nd = not detected. (**B**) Cytotoxicity was assessed by collecting culture supernatants at 48 and 120 hpi and performing a lactate dehydrogenase (LDH) cytotoxicity assay using (+) Glc sample as the spontaneous control. (**C**) Viral to cellular genomes were quantified at the indicated times by quantitative PCR (qPCR). (**D**) Whole cell lysates were collected at the indicated times and analyzed by western blot. A representative blot from three biological replicates is shown. (**E-G**) Viral protein levels were normalized to tubulin levels and quantified relative to (+) Glc at 120 hpi. The graphs display three biological replicates (circles) and one (**D-G**) to three technical replicates. Error bars represent standard deviation (SD). Two-way ANOVA with Tukey’s (**C**) or Šídák’s (**B, E-G**) test were used to determine significance. Statistics were performed on transformed data for **C**. *P* < 0.05, *; *P* < 0.01, **; *P* < 0.001, ***; *P* < 0.0001, ****.

Since glucose supports nucleotide synthesis, we next examined viral genome synthesis from 4-120 hpi using quantitative PCR (qPCR). Viral genome levels were attenuated by 1- to 1.3-log starting at 48 hpi (i.e., at the onset of viral genome synthesis) in glucose-free cultures compared to glucose-replete (**Fig. 1C**). Although viral genomes were significantly decreased from glucose-replete cultures, genomes still increased over time in glucose-free cultures, suggesting low levels of viral genome replication occur independent of glucose.

Glucose is also used to generate several amino acids that support viral protein synthesis. We measured viral protein levels from 4-120 hpi to determine if glucose is necessary to support viral protein production in each of the canonical kinetic classes of lytic replication (i.e., proteins encoded by immediate-early, early, and late viral genes). Immediate-early protein 1 (IE1) and immediate-early protein 2 (IE2) levels during glucose deprivation were similar to glucose-replete conditions up to 48 hpi, suggesting little impact to immediate-early events (**Fig. 1D** and **E; S1A** and **B**). In contrast, early protein UL38 was significantly decreased by 48 hpi (**Fig. 1D** and **F**). Another early protein UL44 was also reduced, but this decrease was only significant at 96 and 120 hpi (**Fig. S1A** and **C**). Late protein pp28 was decreased from 48-120 hpi (**Fig. 1D** and **G**). A second late protein, pp71, was similarly reduced at 96 and 120 hpi in glucose-free cultures (**Fig. S1A** and **D**). A reduction in proteins encoded by late genes is consistent with a loss in viral genome synthesis since the expression of late genes depends on viral genome replication. Taken together, these data indicate that glucose supports early viral protein production and viral genome synthesis as well as subsequent production of late viral proteins. Moreover, our findings suggest that the loss of infectious virus production caused from glucose deprivation is due to defects in early events, and subsequent late steps, of lytic replication.

### Glucose is essential for infectious virus production prior to 48 hpi and inhibition of infectious virus production is reversible with glucose addition

Based on our initial observations that glucose supports early steps in virus replication, we investigated if the effects of glucose deprivation on HCMV replication are time dependent. Attenuated viral genome levels suggest that glucose is required for early replication events prior to, or at the onset of, viral genome replication. We reasoned that glucose removal after viral genome synthesis is initiated would allow early events to proceed as expected but reduce activity in any glucose-dependent steps in the late stages. We infected HFFs with TB40/E-GFP in glucose-replete DMEM, then removed glucose at 1, 24, 48, 72, or 96 hpi and quantified viral titers at 120 hpi. As controls in these experiments, viral titers were determined for glucose-replete or glucose-free cultures. As we previously observed, virus production was not detected in glucose-free conditions when glucose was removed at 1 hpi compared to glucose-replete cultures (**Fig 2A**). Glucose removal at 24 hpi also resulted in undetectable levels of infectious virus. Plaque formation below the limit of quantitation (LoQ) for TCID_50_ was detected when removing glucose at 48 hpi while glucose removal at 72 hpi resulted in measurable virus production that was reduced compared to glucose-replete. Glucose removal at 96 hpi had no effect on viral titers at 120 hpi.

**Fig 2.**
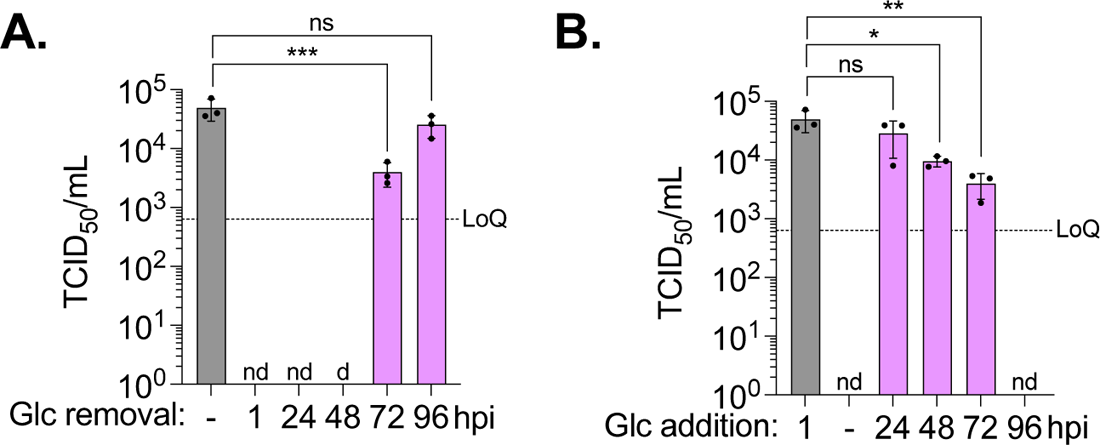
Glucose is required for virus production up to 48 hpi and inhibition of virus production during glucose deprivation is reversible with glucose addition. HFFs were infected with TB40/E-GFP at a MOI 2. At 1 hpi, media was changed to glucose-replete or glucose-free DMEM. (**A**) For glucose removal studies, glucose-replete DMEM was removed, cells were washed with PBS, and media replaced with glucose-free DMEM at the indicated times. (**B**) For glucose addition studies, glucose-free DMEM was removed, cells were washed with PBS, and media replaced with glucose-replete DMEM at the indicated times. Viral titers from culture supernatant were quantified at 120 hpi by TCID_50_. LoQ = 632 TCID_50_/mL; nd = not detected; d = detected but below the LoQ. Experiments from **A** and **B** were performed concurrently using glucose-replete media (grey bar) or glucose-free media (nd) treated at 1 hpi as controls for both. The graphs display three biological replicates (circles) and two technical replicates. Error bars represent SD. One-way ANOVA with Dunnett’s test was used to determine significance. Statistics were performed on transformed data. *P* < 0.05, *; *P* = 0.01, **; *P* < 0.001, ***.

We performed the reciprocal experiment (i.e., time of addition) to determine if the impact of glucose deprivation is reversible. Cultures were glucose-starved at 1 hpi until glucose was added back at 24, 48, 72, or 96 hpi. We found glucose addition at 24 hpi supports virus production similar to glucose-replete whereas virus produced during glucose add back at 48 or 72 hpi was reduced by ∼1-log (**Fig. 2B**). Virus production at 120 hpi was not detected with glucose addition at 96 hpi, potentially due to insufficient time for recovery. These data demonstrate that defects in virus production during glucose loss are reversible with glucose addback. Further, since we observe a defect in early replicative steps and glucose dependence up to 48 hpi, our results suggest that glucose is primarily required for early replication events.

### Uridine and ribose supplement support glucose-independent viral genome synthesis and partially restore infectious virus production

Since glucose is necessary for viral genome replication, we hypothesized metabolites that restore viral genome synthesis in the absence of glucose would promote late viral protein production, and potentially, late replicative steps. Nucleotide synthesis is supported by the conversion of glucose to ribose 5-phosphate via the pentose phosphate pathway (PPP). Uridine has been shown to feed the PPP and restore cancer cell proliferation through the production of ribose 1-phosphate (34). We reasoned that uridine may provide a similar restoration to HCMV genome synthesis in glucose-free cultures. Since DMEM lacks uridine, we tested this idea by feeding uridine to infected cells and measuring viral genome levels. We infected cells as previously described, supplemented with 0-1 mM uridine in glucose-free DMEM at 1 hpi, and measured viral genome levels and virus production at 120 hpi. As we previously observed, viral genomes were reduced in glucose-free cultures compared to glucose-repleted without uridine addition (**Fig. 3A**). Uridine supplement restored viral genomes with an ∼1.5-log increase at 0.5 and 1 mM concentrations compared to glucose-free, non-supplemented cultures. While uridine supplement in glucose-free cultures increased viral genome replication relative to the non-supplemented conditions, viral genomes were still reduced compared to glucose-replete cultures at 120 hpi. Thus, uridine supplement provides a partial rescue of viral genome synthesis. Next, we examined if the partial restoration of viral genome synthesis would lead to production of infectious virus. We found that 1 mM uridine resulted in detectable plaque formation that was below the limit of quantitation (LoQ) for one of two biological replicates (**Fig 3B** and **C**). Further, we found that 0.1 to 1 mM uridine supplement, but not 0 to 0.05 mM, resulted in plaque formation in TCID_50_ cultures despite being below the LoQ (**Fig 3B**). Overall, our observations demonstrate that uridine supplement restores low levels of virus production in glucose-free cultures.

**Fig 3.**
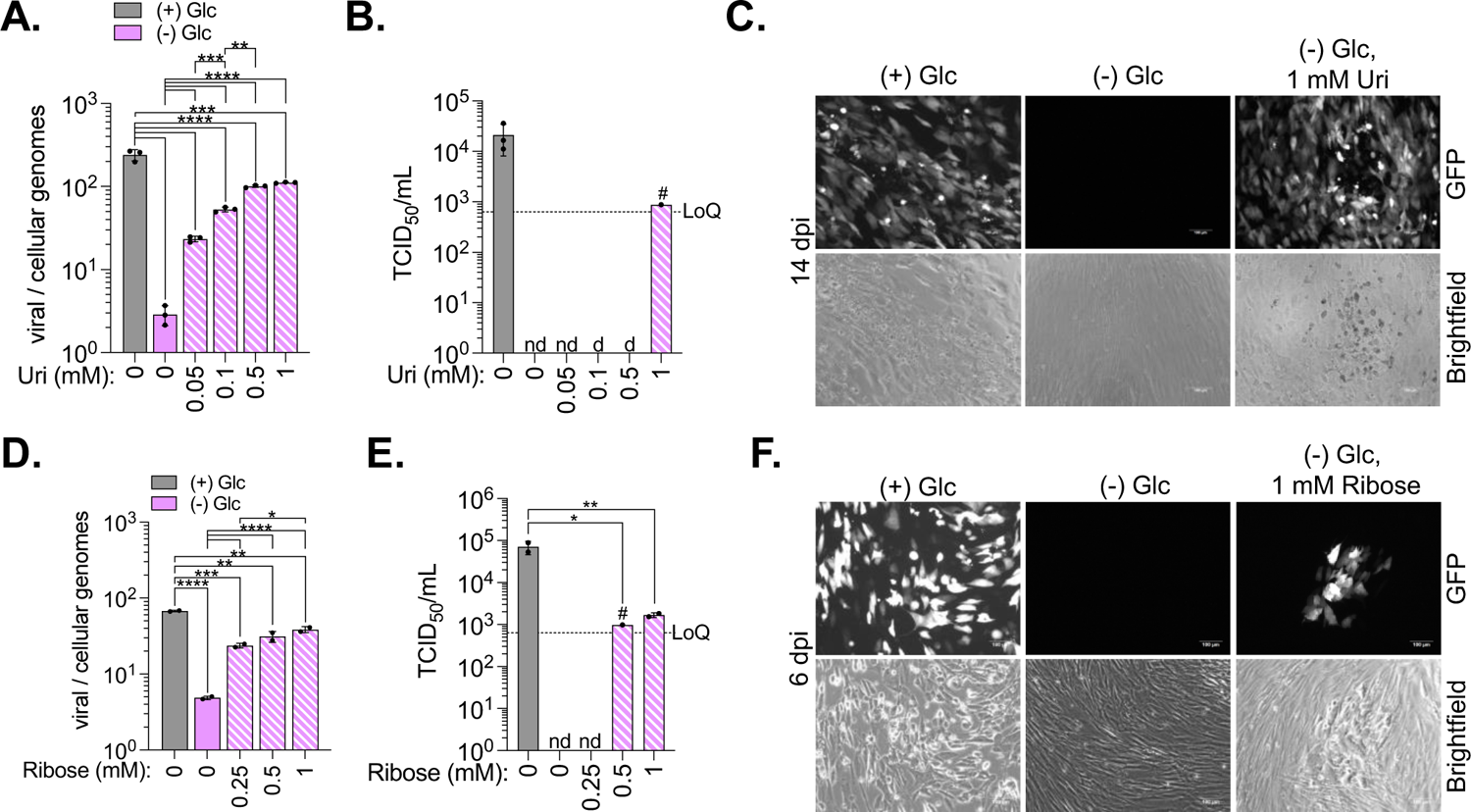
Ribose and uridine supplement of glucose-free cultures partially restores viral genome levels and virus production. (**A-F**) HFFs were infected with TB40/E-GFP at a MOI 2. At 1 hpi, media was changed to glucose-replete or glucose-free DMEM with 0-1 mM uridine (**A-C**) or 0-1 mM ribose (**D-F**). (**A** and **D**) Viral to cellular genomes were quantified by qPCR. (**B** and **E**) Viral titers from culture supernatant at 120 hpi were quantified by TCID_50_. LoQ = 632 TCID_50_/mL; nd = not detected; d = detected but below the LoQ. Pound sign (#) indicates that one of two-three biological replicates had measurable titers. (**C** and **F**) HFFs were infected and treated as described above. At 120 (**C**) or 144 hpi (**F**), culture supernatants were collected, diluted 1:10 in glucose-replete DMEM, and applied to uninfected, subconfluent HFFs. Cells were incubated for 6 (**F**) or 14 d (**C**) and images were taken of GFP-positive plaques. Scale bar = 100 µm. The graphs display two (**D** and **E**) to three biological (circles) and two-three technical replicates. Error bars represent SD. One-way ANOVA with Tukey’s (**D-E**) or Šídák’s (**A**) test were used to determine significance. Statistics were performed on transformed data for **A, D**, and **E**. *P* < 0.05, *; *P* < 0.01, **; *P* < 0.001, ***; *P* < 0.0001, ****.

Uridine restores PPP activity during glucose limitation by replacing glucose as the ribose 5-phosphate donor (34, 35). We hypothesized that if uridine restored viral genome synthesis and low-level virus production via release of ribose, then directly supplementing with ribose would result in a similar rescue. We infected cells, supplemented 0-1 mM ribose in glucose-free DMEM at 1 hpi, and quantified viral genomes and virus production at 120 hpi. Similar to uridine, ribose partially restored viral genome levels with an ∼1-log increase at 0.5 and 1 mM concentrations compared to glucose-free, non-supplemented cultures (**Fig. 3D**). Supplement of 1 mM ribose in glucose-free cultures resulted in quantifiable titers, and 0.5 mM ribose supplement resulted in plaque formation near or below the LoQ (**Fig. 3E**). In other experiments, viral titers for 1 mM ribose supplemented, glucose-free cultures were under the LoQ with detectable plaque formation (**Fig. 3F; S2A**). We confirmed ribose restoration of virus production using a different commercially available ribose stock. Ribose supplement consistently produced infectious virus as evidence by plaque formation but was below the LoQ for three of four biological replicates, confirming that our observation is independent of the company supplying ribose (**Fig. S2B**). These results demonstrate that ribose supplement consistently restores virus production but at levels that are challenging to quantifiably measure.

### Ribose supplement partially restores viral genome synthesis, viral protein levels, and released viral particle levels

Because ribose restored viral genome synthesis during glucose deprivation, we next measured cytotoxicity to determine if ribose supplement could rescue cell viability in mock-infected primary fibroblasts. HFFs were infected or mock-infected and at 1 hpi media was replaced with glucose-replete or glucose-free with 1 mM ribose. At 48 and 120 hpi, culture supernatants were collected for lactate dehydrogenase cytotoxicity assay. At 48 hpi, cells in all conditions maintained high viability in both mock and HCMV-infected cultures (**Fig. 4A**). At 120 hpi, mock-infected, glucose-replete, ribose supplemented cultures maintained high cell viability but mock-infected, glucose-free, ribose supplemented cell viability decreased to 76%, indicating ribose is not sufficient to protect HFFs from glucose deprivation-induced cytotoxicity. HCMV-infected, glucose-free, ribose supplement cultures maintained 95% cell viability while HCMV-infected, glucose-replete, ribose supplemented cultures decreased to 86%, potentially due to cell death from HCMV replication similar to HCMV-infected, glucose-replete cultures not supplemented with ribose (**Fig. 1B**). Our findings suggest that ribose supplement does not impact cell viability in glucose deprived mock or HCMV-infected cells.

**Fig 4.**
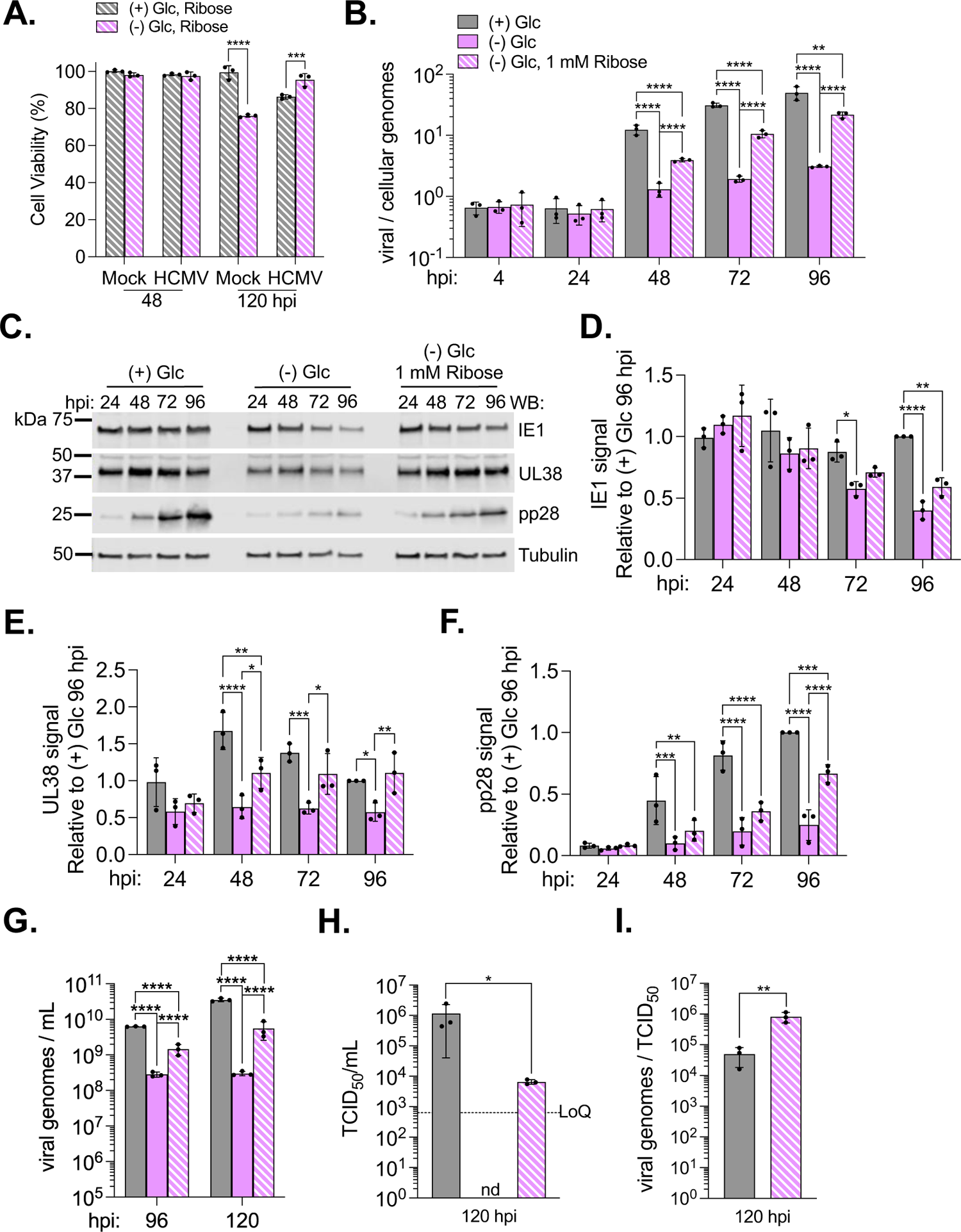
Ribose partially restores viral genome synthesis, viral protein levels, and released particle levels. HFFs were infected with TB40/E-GFP at a MOI 2. At 1 hpi, media was changed to glucose-replete or glucose-free supplemented with 1 mM ribose (**A**) or glucose-replete, glucose-free, or glucose-free with 1 mM ribose supplement (**B-I**). (**A**) Cell viability was measured by collecting culture supernatants at 48 and 120 hpi and performing a lactate dehydrogenase (LDH) cytotoxicity assay using (+) Glc as the spontaneous control. (**B**) Viral to cellular genomes were quantified by qPCR at the indicated times. (**C**) Whole cell lysates were collected at the indicated times and analyzed by western blot. A representative blot from three biological replicates is shown. (**D-F**) Viral protein levels were normalized to tubulin levels and quantified relative to (+) Glc at 96 hpi. (**G**). At 96 and 120 hpi, 100 µl of culture supernatants were collected, treated with DNase and released particle genomes were quantified by qPCR. (**H**) Viral titers were quantified by TCID_50_ using the same culture supernatant from **G** at 120 hpi. LoQ = 632 TCID_50_/mL; nd = not detected (**I**) The ratio of total particles released to infectious particles released was determined using data from **H** and **G**. The graphs display three biological (circles) and one (**C-F**)-three technical replicates. Error bars represent SD. Two-way ANOVA with Tukey’s (**A, D-F, G**) or Šídák’s (**B**) test or paired t-test (**H** and **I**) were used to determine significance. Statistics were performed on transformed data for **A** and **G-I**. *P* < 0.05, *; *P* < 0.01, **; *P* < 0.001, ***; *P* < 0.0001, ****.

We continued to supplement with ribose directly to better understand how HCMV might use compensatory nutrients to replicate during glucose deprivation. As ribose partially restores viral genome synthesis by 120 hpi, we next investigated if ribose supplement rescues viral genome synthesis. We measured viral genome levels from 4-96 hpi in glucose-replete or glucose-free cultures with or without 1 mM ribose. Ribose supplement significantly increased viral genome levels relative to glucose-free, non-supplemented cultures beginning at 48 hpi (**Fig. 4B**). Again, viral genome levels in glucose-free, ribose supplement were reduced compared to glucose-replete, indicating ribose partially restores viral genome synthesis in glucose limiting conditions.

Because ribose partially restores viral genome synthesis, we reasoned that viral protein production would also be restored in supplemented cultures without glucose. We measured viral proteins levels from 4-96 hpi in glucose-replete or glucose-free DMEM with or without ribose. For immediate-early proteins, we observed that IE1 and IE2 levels remained unchanged in glucose-free, ribose supplemented cultures compared to glucose-replete until late stages of virus replication when IE1 levels decreased (**Fig. 4C** and **D; S3A** and **B**). For early proteins, ribose supplement partially rescued UL38 levels by 48 hpi relative to glucose-free, non-supplemented conditions and provided a full restoration at 96 hpi relative to glucose-replete conditions (**Fig. 4C** and **E**). UL44 levels remained similar compared to glucose-free conditions until 96 hpi when levels were increased compared to glucose-free cultures yet still decreased from glucose-replete (**Fig. S3A** and **C**). Finally, late viral proteins pp28 and pp71 levels in ribose supplement cultures were partially restored (**Fig. 4C** and **F, S3A** and **D**). These findings demonstrate that ribose supports low levels of glucose independent HCMV production via partial restoration of viral genome synthesis and rescue of viral protein levels.

Since ribose supplement in glucose limiting conditions partially supported viral genome synthesis, early and late protein levels, and a low level of infectious virus production, we investigated if ribose supplement was able to restore late events leading to release of virus particles. To determine if viral particles are released during glucose deprivation with or without ribose, we infected and treated HFFs as previously described, collected and DNase treated culture supernatant, and quantified protected viral genomes as a measurement of released viral particles. Viral particles were decreased during glucose deprivation by ∼1- and 2-log at 96 and 120, respectively, and ribose supplement increased levels by ∼1-log compared to glucose-free at both timepoints (**Fig. 4G**). We quantified the ratio of total released particle levels (**Fig. 4G**) to infectious particle levels (**Fig. 4H**) as a measure of particle infectivity (**Fig. 4I**). The ratio of total released particles to infectious particles was increased by ∼1-log in ribose supplemented, glucose-free cultures compared to glucose-replete, demonstrating that particles released in ribose supplemented conditions are less infectious than those made in glucose-replete conditions (**Fig. 4I**). These results indicate that ribose supports particle production during glucose deprivation, but these particles still lack components required for infectivity.

### Ribose supplement partially restores glycosylated gB levels in glucose-free cultures

Viral glycoproteins are responsible for particle binding, fusion, and entry events into host cells. Based on our observation that ribose supplement yields released particles that are deficient in their ability to establish infection in cells, we considered that ribose may not restore the role of glucose in the formation of functional glycoproteins. Glucose contributes to glycosylation through UDP-glucose or the synthesis of UDP-GlcNAc via the hexosamine pathway. Loss of UDP-sugars result in decreased viral protein glycosylation and infectious virus production (23, 24). To determine if viral protein glycosylation is lost during glucose deprivation, we examined a representative HCMV glycoprotein, glycoprotein B (gB). In glucose-replete cultures, we observed a band just below 150 kDa that increased intensity over time (**Fig. 5A** and **B**). The ∼150 kDa band was not evident in glucose-free cultures but darkening the image revealed the 150 kDa band at 24 hpi. The band was absent at later timepoints. As the 150 kDa band is present at 24 h in all conditions, it is likely residual glycosylated gB from inoculating viral particles. Darkening the image also revealed another band around 100 kDa in glucose-free conditions from 48-96 hpi. In contrast, ribose supplemented, glucose-free cultures displayed a smear from ∼150 to ∼100 kDa that increased over time with the strongest intensity at 96 hpi. Quantification of the ∼150 kDa portion of the smear suggests that ribose partially restores some level of this band relative to glucose-free cultures (**Fig. 5B**). The ∼150 kDa band likely corresponds to glycosylated gB, while the band near 100 kDa band may be unglycosylated gB that runs at an expected size of 105 kDa (36). To confirm this observation, we collected whole cell lysates from glucose-replete cultures at 96 hpi and hydrolyzed N-linked glycans using PNGase F enzyme.

**Fig 5.**
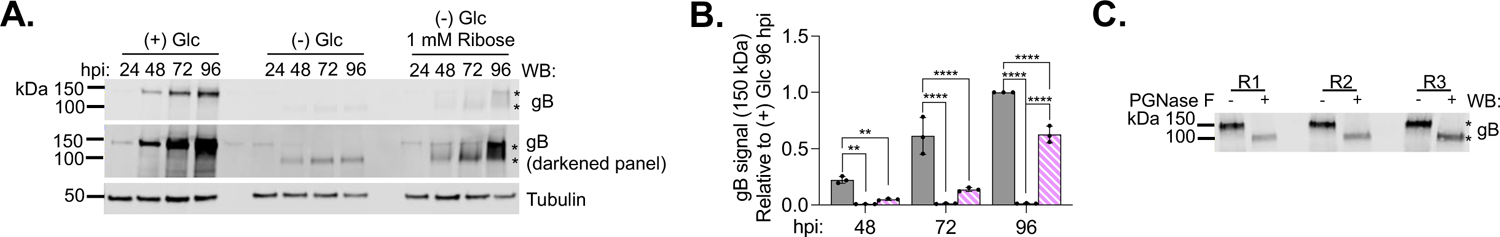
Ribose partially restores gB glycosylation during glucose deprivation. (**A**) Whole cell lysates were collected at the indicated times and analyzed by western blot using antibodies to gB and tubulin. Asterisks (*) indicate ∼150 kDa (upper) and ∼105 kDa (lower) gB bands. A representative blot from three biological replicates is shown. (**B**) The ∼150 kDa gB levels were normalized to tubulin levels and quantified relative to (+) Glc at 96 hpi. (**C**) HFFs were infected as described above and cultured in glucose-replete media. Whole cell lysates were collected, total protein quantified, and 15 μg of protein incubated with PNGase F to hydrolyze N-linked oligosaccharides. Lysates were analyzed by western blot. A blot with three biological replicates is shown. The graphs display three biological replicates (circles). Error bars represent SD. Two-way ANOVA with Tukey’s test was used to determine significance. *P* < 0.01, **; *P* < 0.0001, ****.

The ∼150 kDa band shifted to 105 kDa following PNGase F treatment (**Fig. 5C**), indicating that this upper band is glycosylated gB and the 105 kDa band is unglycosylated gB. These results suggest that ribose partially restores glycosylated gB levels by 96 hpi during glucose deprivation.

### Metabolites that support viral genome synthesis and glycosylation during glucose deprivation restore low levels of infectious virus production

Since glycosylation of viral receptors is required for particle infectivity (37), we hypothesized that restoration of protein glycosylation with rescue of viral genomes would restore particle infectivity in glucose-free cultures. To this end, we supplemented glucose-free cultures with ribose and/or UDP-N-acetylglucosamine (UDP-GlcNAc), the substrate for both O- and N-linked glycosylation reactions. HFFs were infected as previously described. At 1 hpi, media was changed to glucose-replete, glucose-free, or glucose free supplemented with 1 mM ribose, 100 µM UDP-GlcNAc, or both metabolites (dual supplement). Whole cell lysates were collected at 120 and 144 hpi and protein glycosylation was evaluated by blotting for gB. In glucose-replete conditions, the ∼150 kDa gB band was observed, while only unglycosylated gB was observed in glucose-free conditions (**Fig. 6A**) Ribose supplement restored glycosylation activity as evidenced by a prominent ∼150 kDa band at 120 and 144 hpi. Glycosylated gB levels during ribose or dual supplement were fully restored to glucose-replete levels as were both late proteins pp28 and pp71 (**Fig. 6A-C; S4A** and **B**). UDP-GlcNAc supplement resulted in upper and lower gB bands present at both timepoints (**Fig. 6A**) and glycosylated gB was significantly reduced compared to either glucose-replete or ribose supplemented, glucose-free cultures (**Fig. 6A** and **B**). A possible explanation for the observation that ribose restores gB glycosylation to a greater extent than UDP-GlcNAc is that gB is a late protein and expression of late proteins is greater in ribose supplement than UDP-GlcNAc supplement. We examined late protein pp28 and found that UDP-GlcNAc increased pp28 levels compared to glucose-free with fully restored levels by 144hpi (**Fig. 6A** and **C**). Additionally, pp71 levels were fully restored by 120 hpi in UDP-GlcNAc supplemented conditions (**Fig. S4A** and **B**). Together, these data suggest UDP-GlcNAc supplement impacts late viral protein levels during glucose deprivation yet only marginally restores glycosylation.

**Fig 6.**
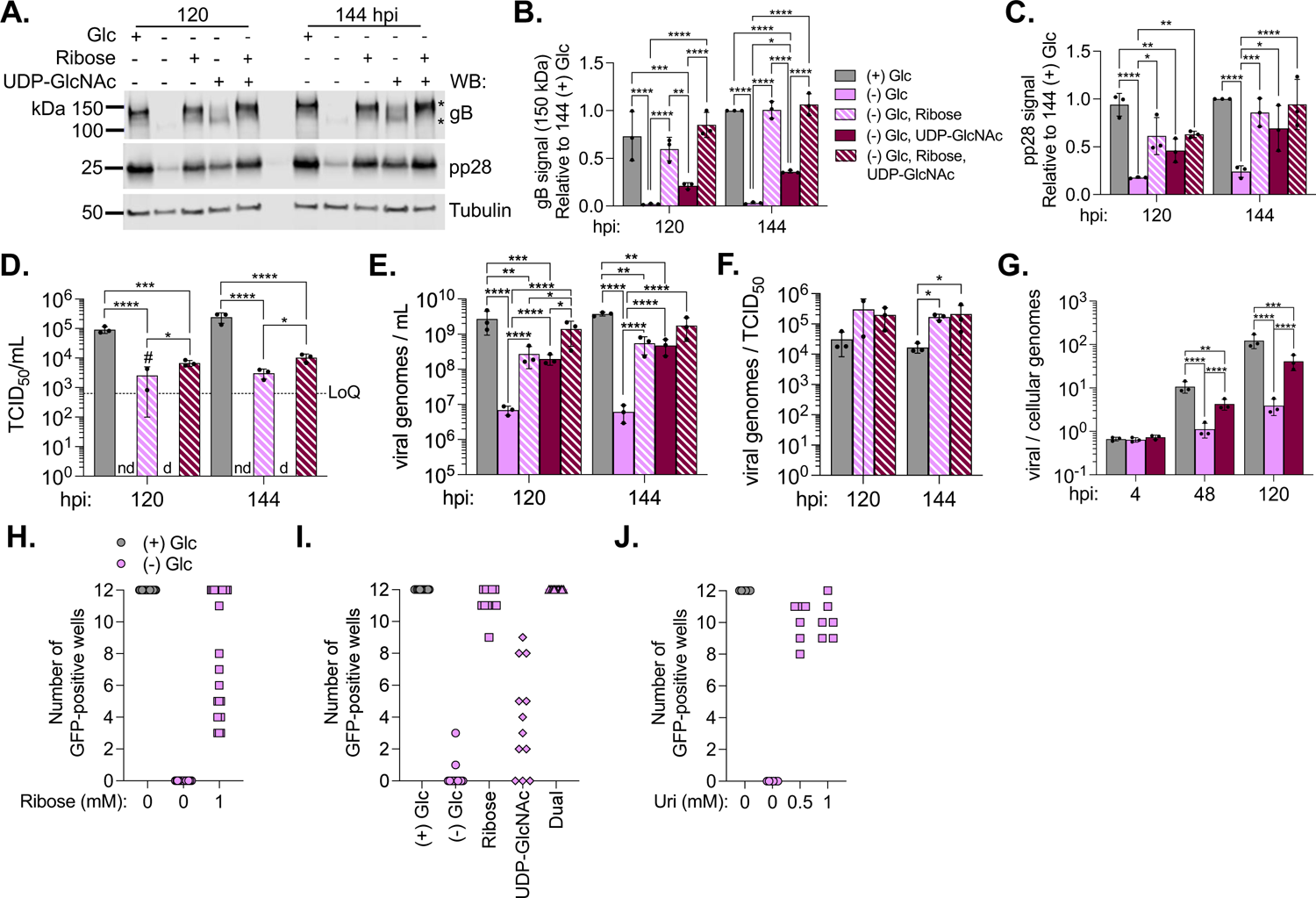
Metabolites that support viral genome synthesis and glycosylation restore virus production during glucose deprivation. HFFs were infected with TB40/E-GFP at a MOI 2. At 1 hpi, media was changed to glucose-replete, glucose-free, or glucose-free DMEM supplemented with 1 mM ribose, 100 µM UDP-N-acetylglucosamine (UDP-GlcNAc), or both. (**A**) Whole cell lysates were collected at 120 and 144 hpi and analyzed by western blot. Asterisks (*) indicate 150 kDa (upper) and 105 kDa (lower) gB bands. A representative blot from three biological replicates is shown. (**B** and **C**) The levels of gB and pp28 were normalized to tubulin levels and quantified relative to (+) Glc at 144 hpi. (**D-F**) At 120 and 144 hpi, viral titers were quantified from culture supernatant by TCID_50_ (**D**) and released particle genomes from 100 µl of culture supernatant were quantified by qPCR (**E**). (**F**) The ratio of total particles released to infectious particles released was determined using data from **D** and **E**. LoQ = 632 TCID_50_/mL: nd = not detected; d = detected but below the LoQ. Pound sign (#) indicates that two of three biological replicates had measurable titers. (**G**) HFFs were infected with TB40/E-GFP at a MOI 2. At 1 hpi, media was changed to glucose-replete, glucose-free, or glucose-free DMEM supplemented with 100 µM UDP-GlcNAc. Viral to cellular genomes were quantified by qPCR at the indicated timepoints. (**H-J**) Graphs display the number of GFP-positive wells out of 12 wells of a 96-well dish that were infected with a 1:10 dilution of culture supernatant from 120 hpi from experiments conducted using 1 mM ribose (n=11) (**H**), 1 mM ribose, 100 µM UDP-GlcNAc, or dual ribose and UDP-GlcNAc (n=6) (**I**), or 0.5 and 1 mM uridine (n=3) (**J**) supplement. Each dot represents a technical replicate from the total biological replicates indicated above. Graphs **B-G** display three biological replicates (circles) and one (**A-C**) to three technical replicates. Error bars represent SD. Two-way ANOVA with Tukey’s test was used to determine significance. Statistics were performed on transformed data for **D** and **E**. *P* < 0.05, *; *P* < 0.01, **; *P* < 0.001, ***; *P* < 0.0001, ****.

We next examined viral titers, total released particles, and particle infectivity during supplementation. Again, no infectious virus production was detected in glucose-free cultures, and ribose partially restored virus production in glucose-free cultures to measurable levels at 120 and 144 hpi (**Fig. 6D**). UDP-GlcNAc supplement alone was sufficient to restore low levels of infectious virus production below the LoQ as plaque formation was evident on the TCID_50_ plates. Dual supplement increased virus production by ∼0.5-log compared to ribose supplement alone. However, virus production was not recovered to the levels observed in glucose-replete conditions.

Ribose and UDP-GlcNAc supplemented individually partially restored total released particles compared to glucose-free cultures (**Fig. 6E**). Dual supplement of both ribose and UDP-GlcNAc fully restored viral particle levels. Particle infectivity was calculated for conditions with measurable titers. Both ribose and dual supplement cultures had a ∼1-log increase in noninfectious particles at 144 hpi compared to glucose-replete (**Fig 6F**), suggesting these particles still lack components require for infectivity despite restoration of viral protein glycosylation.

Because UDP-GlcNAc restored both late viral protein and released viral particles during glucose deprivation, we hypothesized viral genome synthesis was also restored. We infected HFFs as previously described and treated with glucose-replete, glucose-free, or glucose free with 100 µM UDP-GlcNAc. We collected timepoints from 4-120 hpi and quantified viral genome synthesis using qPCR. UDP-GlcNac partially restored viral genomes levels at 48 hpi and 120 hpi (**Fig. 6G**). Together with the inefficient restoration of gB glycosylaton during UDP-GlcNAc supplement, these data suggest UDP-GlcNAc is prioritized for other metabolic processes aside from protein glycosylation such as viral genome synthesis.

Ribose, UDP-GlcNAc, and uridine can feed anabolic pathways that branch from upper glycolysis. Our data demonstrate that these metabolites partially restore viral genome synthesis and support low virus production in glucose-free cultures. To better display this consistent low level of virus production, we collected the supernatant from infected cells grown in supplemented and non-supplemented conditions, diluted it 1:10 and inoculated 12 wells of uninfected cells in 96-well plates. We then counted the number of wells that contain at least one GFP-positive virus plaque. We used this approach as a semi-quantitative measure of virus produced below the LoQ for our standard TCID50 assay. As expected, in glucose-replete conditions, all 12 wells were positive (**Fig. 5H-J**). For glucose-free cultures, two plates from a total of 41 plates had one and three GFP-positive wells, respectively. In contrast, ribose, dual ribose and UDP-GlcNAc, and uridine supplement of glucose-free cultures consistently resulted in plaque formation that did not always reach the level where all 12 wells were positive (**Fig 3C** and **F; 5H-J**). For UDP-GlcNAc supplement, nine out of 12 plates had detectable GFP-positive plaques (**5I**). Our observations demonstrate that compensatory nutrients can support low levels of HCMV production in glucose-free conditions.

We next asked if restoration of lower glycolysis in glucose-free cultures would rescue HCMV replication. To this end, we supplemented glucose-free cultures with pyruvate, the final product of glycolysis. We infected HFFs, supplemented with 0-4 mM pyruvate in glucose-free DMEM, and measured viral genome levels and virus production at 120 hpi. Higher concentrations of pyruvate supplement in glucose-free cultures negligibly increased viral genome levels compared to glucose-free, non-supplemented cultures (**Fig. 7A**). All pyruvate concentrations tested failed to produce detectable plaque formation on titer plates (**Fig. 7B**), indicating that pyruvate is not sufficient to restore viral replication. Overall, our findings demonstrate that metabolites that feed upper glycolytic branches such as the PPP are sufficient to support low levels of glucose independent virus production, but supplement of lower glycolysis cannot restore virus production.

**Fig 7.**
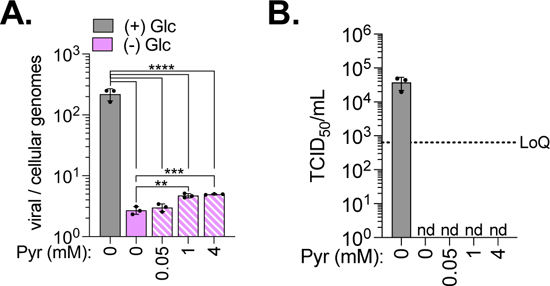
Pyruvate does not restore viral genome synthesis or virus production in glucose-free cultures. HFFs were infected with TB40/E-GFP at a MOI 2. At 1 hpi, media was changed to glucose-replete or glucose-free DMEM with 0-4 mM pyruvate. (**A)** Viral to cellular genomes were quantified by qPCR. (**B**) Viral titers from culture supernatant at 120 hpi were quantified by TCID_50_. LoQ = 632 TCID_50_/mL; nd = not detected. Šídák’s test was used to determine significance. Statistics were performed on transformed data. *P* < 0.01, **; *P* < 0.001, ***; *P* < 0.0001, ****.

## DISCUSSION

HCMV relies on manipulation of host metabolism to replicate and does so successfully in various human tissue with different nutrient environments. Most HCMV metabolic studies are completed in high nutrient cell culture media that contain supraphysiological levels of some nutrients while lacking other metabolites found in human sera (7–10, 19, 21, 22, 24–26, 32, 38, 39). While these studies have advance our understanding by identifying metabolic alterations caused by virus infection, they lead to two questions: 1) do nutrients found in human sera but absent from typical cell culture media compensate and contribute to HCMV replication when other metabolites are limited? 2) do supraphysiological nutrient concentrations mask the contribution of other nutrients to HCMV replication? We address these questions using a glucose independent model of HCMV infection to demonstrate that viral genome synthesis requires glucose but metabolites that support upper glycolytic branches compensate to restore viral genome replication and low levels of virus production. Our data suggests that in limited glucose conditions, ribose feeds nucleotide synthesis and non-oxidative PPP to support downstream replication processes such as viral genome synthesis, viral protein production, and protein glycosylation, resulting in low but notably increased virus production. Overall, our findings indicate that HCMV successfully replicates in glucose limited environments by nutrient compensation that feeds metabolic pathways necessary for replication.

Uridine supplement partially restores viral genome synthesis in glucose-free cultures (**Fig. 3A**). Uridine-derived ribose 1-phosphate could restore viral genome synthesis through its conversion to ribose 5-phosphate and then phosphoribosyl diphosphate (PRPP) to support both purine and pyrimidine synthesis (**Fig. 8**). In support of this possibility, direct ribose supplement restored viral genome synthesis to a similar level as uridine supplement (**Fig. 3D**). Similar to our observations, studies in cancer cell lines demonstrated that uridine supports glucose independent cell proliferation and protects from glucose deprivation-induced cell death through uridine phosphorylase activity and ribose 1-phosphate release (34, 35, 40–44).

**Fig 8.**
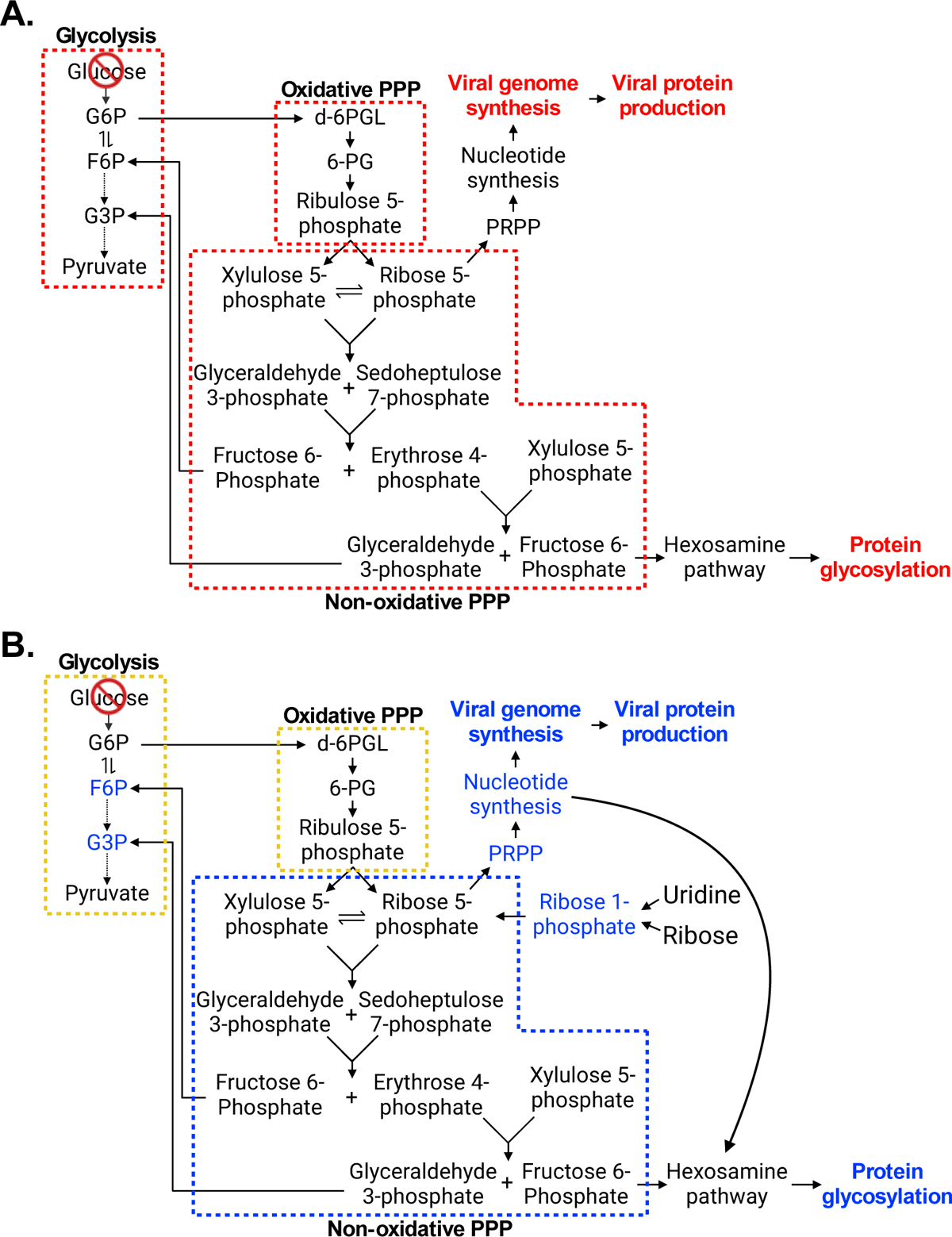
Model of metabolite regulation during glucose deprivation. Schematic of abbreviated glycolysis and the pentose phosphate pathway during glucose deprivation (**A**) and uridine or ribose supplement during glucose deprivation (**B**). (**A**) Metabolic pathways boxed in red are likely inhibited during glucose deprivation. Processes in red were decreased in glucose-free conditions. (**B**) Metabolic pathways or metabolites in blue are likely restored in glucose limiting conditions when supplemented with ribose or uridine, while pathways boxed in yellow are potentially restored. Processes in blue were increased in ribose or uridine supplemented, glucose-free cultures compared to glucose-free.

Our data supports a model in which uridine-derived ribose supports overall nucleotide synthesis via conversion to PRPP in glucose limiting cultures (**Fig. 8**). Compensation by uridine-derived ribose during glucose deprivation could be a mechanism for how HCMV successfully replicates in glucose limiting environments. When supplementing with ribose directly, we found that in addition to restoring viral genome synthesis (**Fig. 3D, 4B**), ribose also rescues glycosylated gB levels by 120 hpi (**Fig. 6A** and **B**). This restoration could occur through anabolic reactions that contribute metabolites for UDP-GlcNAc synthesis through the hexosamine pathway. Ribose rescue of nucleotide synthesis would result in UTP availability for the last step of UDP-GlcNAc synthesis (**Fig. 8**). Ribose could also feed the non-oxidative branch of the PPP that yields fructose 6-phosphate, which is the first substrate in the hexosamine pathway. In support of this potential mechanism, ribose 1-phosphate feeding non-oxidative PPP was found necessary for cancer cell proliferation during glucose starvation (34). Based on these studies, we conclude that ribose restoration of gB glycosylation likely results from ribose feeding non-oxidative PPP that produces fructose 6-phosphate to support UDP-GlcNAc synthesis and glycosylation during glucose limitation.

Ribose supporting non-oxidative PPP would also support lower glycolysis through formation of glyceraldehyde 3-phosphate, or potentially, support both upper and lower glycolysis by feedback to glucose 6-phosphate via the enzyme glucose 6-phosphate isomerase that converts fructose 6-phosphate to glucose 6-phosphate (**Fig. 8**) (34). Our data demonstrate that pyruvate supplement failed to restore viral genomes levels or virus production (**Fig. 7**), indicating that restoring only lower glycolysis is not sufficient to restore virus production in glucose-free conditions. Thus, metabolites that support upper glycolytic branches are necessary for HCMV virus production in glucose limiting conditions (**Fig. 3B** and **C, E** and **F; 6H-J**). These results indicate that a major role of glucose during HCMV replication is support of viral genome synthesis and protein glycosylation and these processes are rate limiting steps in glucose independent HCMV replication.

Our data indicate that HCMV offers protection against glucose deprivation-induced cytotoxicity as HCMV-infected, glucose-free cultures have increased cell viability compared to mock-infected cells by 120 hpi (**Fig. 1B**). These results are in agreement with prior studies conducted in HFFs during glucose starvation by Chambers, et al. (19). In contrast, a recent study by Raymonda, et al. (20) showed that mock-infected MRC5 fibroblasts starved of glucose remain viable, while HCMV-infected cells have decreased viability by 48 hpi. There were several differences noted between the Chambers and Raymonda studies such as the presence or absence of FBS, cell type, and assay times. Our conditions of fully confluent, serum-starved cells and a brief infection period (1-2 h) were similar to Raymonda, et al., suggesting these conditions are unlikely to be contributing to death of HCMV-infected cells during glucose starvation that was not observed in our study or Chambers, et al. These conflicting results could be due to differences in the fibroblast cells used: HFF versus embryonic lung MRC5 fibroblast cells. Additionally, Raymonda, et al., used the lab-adapted AD169 strain whereas Chambers, et al. used Towne and we used TB40/E. It is conceivable that there are differences between the response of primary human fibroblast lines to glucose deprivation independent of infection or differences in interactions between HCMV TB40/E or Towne and fibroblasts compared to AD169 and fibroblasts in glucose limiting conditions.

Our findings reveal that the presence or absence of certain nutrients impacts phenotypes observed in glucose-free conditions. In rich conditions, glucose is used to supply nucleotide synthesis, glycosylation, lipid synthesis (7–10, 17, 18, 22–26). In this study, we demonstrate that uridine and ribose can be used as an alternative nutrient source for nucleotide synthesis and glycosylation. It is currently unknown if HCMV infection can promote lipid synthesis in glucose limiting conditions. Raymonda, et al., suggest that expression of HCMV UL38 protein in uninfected cells promotes fatty acid synthesis during glucose deprivation (20). It is unknown if UL38 has a similar function during HCMV infection of HFF fibroblasts. Our future studies will focus on regulation of fatty acid and lipid synthesis in nutrient limited environments during HCMV replication to define if HCMV can promote lipid synthesis independent of glucose and if alternative nutrients are prioritized for lipid metabolism during infection.

While our observations show that uridine compensates for glucose starvation, uridine could also contribute to HCMV replication in physiological glucose environments. Munger et al. (8) demonstrated that in high glucose cultures uridine is decreased by ∼4-fold during HCMV replication compared to mock-infected cells. While the authors attributed this decrease to only nucleotide synthesis, it is plausible that uridine contributes ribose for non-oxidative PPP, and subsequently, the hexosamine pathway when glucose is at physiological or sub-physiological concentrations and supraphysiological glucose levels mask the contribution of uridine to this process.

When evaluating the reduction in glycosylation in the absence of glucose, we found that UDP-GlcNAc alone restored viral genome synthesis, late viral protein levels, and virus production with only marginally increased gB glycosylation (**Fig. 6A-D, G** and **I**). We anticipated that UDP-GlcNAc supplement would restore glycosylation of the lttle gB produced in glucose-free cultures (**Fig. 5A** and **6A**) though be unable to rescue late viral protein levels or virus production. UDP-GlcNAc could contribute to viral genome synthesis through UDP loss after glycosylation, which could then be converted to UTP or metabolized to ribose through salvage pathways. However, UDP-GlcNAc was not sufficient to restore glycosylated gB levels even by 144 hpi, suggesting glycosylation may not be the primary role of this metabolite in glucose-free conditions. A possible explanation for the level of glycosylation observed following UDP-GlcNAc supplement is that the nutrient is prioritized for other metabolic reactions during viral replication in limited glucose conditions.

HCMV-infected cells demand glucose for optimal virus production in culture as evidenced by increased glucose uptake and glycolytic flux (7–10, 21, 22). We demonstrate that metabolites that feed upper glycolytic branches compensate for essential roles of glucose to support viral genome synthesis and promote low, consistent levels of virus production. Low, yet persistent, replication of HCMV in the body may have importance in the virus’s ability to spread and cause disease in immunosuppressed individuals. Our results suggest that HCMV takes advantage of alternative metabolite use in nutrient limited conditions to support low levels of virus replication.

## MATERIALS AND METHODS

### Cell culture and virus

Human foreskin fibroblasts (HFFs) were cultured in Dulbecco’s modified Eagle Medium (DMEM) containing 10% fetal bovine serum (FBS), 10 mM HEPES, and penicillin/streptomycin (P/S). Unless otherwise noted, all experiments were performed in 6-well dishes. Prior to infection, cells were held at full confluency for 72 h in full growth medium and then another 24 h prior to infection under serum free (SF) conditions (DMEM, 10 mM HEPES, P/S). Cells were infected at a multiplicity of infection (MOI) 2 infectious units per cell (IU/cell) using HCMV strain TB40/E encoding free GFP (TB40/E-GFP) for 1 h, washed with PBS, and new media added. Mock-infected cells were treated with inoculum that did not contain virus. Unless otherwise noted, media was replaced at 48 hpi to maintain nutrient levels. Cell viability was measured using CyQuant lactate dehydrogenase (LDH) cytotoxicity assay (ThermoFisher Scientific) in a 96-well plate.

Viral stocks were produced by propagating BAC-derived virus in fibroblasts. Virus from culture supernatant were concentrated by pelleting through a sorbitol (20% sorbitol, 50 mM Tris pH 7.2, 1 mM MgCl_2_) cushion at 20,000 rpm for 80 min using ultracentrifugation and resuspension in SF DMEM containing 10 mM HEPES and P/S. Stocks and experimental samples were titered using a 50% tissue culture infectious dose (TCID_50_) assay beginning with a 1:10 dilution of the stock or experimental sample and counting GFP-positive wells at 2 weeks post infection. If there were no GFP-positive wells in the first row of the titer plate, the sample was considered not detected (nd). If the number of GFP-positive wells in the first row was below 12, the sample was detected (d) but below the LoQ.

### Chemical reagents

DMEM without glucose was either purchased (Gibco) or prepared using DMEM powder (Gibco) and adding 4 mM glutamine and 44 mM sodium bicarbonate. DMEM containing glucose was either purchased (Gibco) or prepared in a similar manner with the addition of 25 mM glucose. Ribose (100 mM; Millipore or ThermoFisher Scientific), Uridine (100 mM; Millipore), Pyruvate (100 mM; Millipore), and UDP-GlcNAc (77 mM; Millipore) stocks were maintained in glucose-free, SF DMEM, 10mM HEPES, P/S at 4°C after filter sterilizing.

### DNA quantification

Quantitative PCR (qPCR) was used to assess whole cell DNA and viral particle genome levels. For whole cell DNA, cells were collected by trypsinization and DNA was isolated using Zymo Quick DNA mini-Prep kit (ThermoFisher Scientific). For viral particle genomes, 100 µl of culture supernatant was collected and treated with 2 µl (5 U) DNase solution I (ThermoFisher Scientific) at 37°C for 1 h. Protected viral genomes were isolated using Zymo Quick DNA mini-Prep kit. Viral and cellular genome levels were measured using primers to HCMV UL123 (5’-GCCTTCCCTAAGACCACCAAT-3’ and 5’-ATTTTCTGGGCATAAGCCATAATC-3’) and cellular actin (5’-TCCTCCTGAGCGCAAGTACTC-3’ and 5’-CGGACTCGTCATACTCCTGCTT-3’). Quantities were determined using an absolute standard curve that was prepared using a BAC containing the genome sequence for HCMV strain FIX with the cellular actin gene (45). qPCR was completed using PowerUp SYBR Green Master Mix (ThermoFisher Scientific) and QuantStudio 3 Real Time PCR system.

### Protein analysis

Protein levels were analyzed by western blot. Whole cell lysates were collected by in-well lysis and scraping. For hydrolysis of N-linked glycans, total protein was quantified using Pierce BCA Protein Assay Kit (ThermoFisher Scientific), and the PNGase F assay (New England Biolabs) was completed following the manufacturer’s instructions using 15 µg of protein. Proteins were resolved by SDS-PAGE using Mini-Protean any kD or 4-20% gradient gels (BioRad) and transferred to nitrocellulose membrane (LI-COR). Membranes were blocked in 5% milk in tris buffered saline with 0.1% Tween 20 (TBST). Primary antibodies were diluted in 1% milk in TBST and incubated with rocking for 1 h at room temperature or overnight at 4°C. Secondary antibodies were diluted in 5% milk in TBST and incubated for 1 h with rocking in the dark at room temperature. Images were taken on a LI-COR Odessey CLx imager, and quantitation was performed using Image Studio Lite software.

The following primary antibodies were used for western blot analysis: Mouse-anti UL44 (virusys; 1:2,500), rabbit-anti tubulin (Proteintech; 1:1,000), mouse-anti tubulin (Proteintech; 1:1,000), mouse anti-IE1 (clone 1B12; 1:100), mouse anti-IE2 (clone 3H4; 1:100), mouse anti-UL38 (clone 8D6; 1:100), mouse anti-pp28 (clone 10B4-29; 1:100), mouse anti-pp71 (clone 2H10-0; 1:100), and mouse anti-gB (clone 27-156; 1:50). All HCMV antibodies except for anti-UL44 and anti-gB were gifts from Dr. Thomas Shenk (Princeton University). The anti-gB was provided by Dr. William Britt (University of Alabama at Birmingham) (46). Goat anti-mouse DyLight 800 (1:10,000) or goat anti-rabbit DyLight 680 (1:10,000) were used for secondary antibodies.

### Statistics

Graphs were designed and statistical testing were performed using GraphPad Prism. The appropriate statistical tests for each experiment are noted in the figure legends. Model in Fig 8 was created with Biorender.com.

## AUTHOR CONTRIBUTIONS

R.L.M. and J.G.P. conceptualized and designed experiments; R.L.M performed experiments and analyzed the data. R.L.M. and J.G.P. interpreted the results; R.L.M. drafted the manuscript; R.L.M. and J.G.P. reviewed and edited the manuscript. All authors have read and approved the final manuscript.

## ACKNOWLEDGMENTS

We thank Felicia Goodrum and Jim Alwine for helpful feedback. We thank members of the Purdy and Goodrum labs for their input on the project. This project was supported by the National Institute of Allergy and Infectious Disease division of the National Institute of Health under award numbers R01AI162671, R01AI155539 (J.G.P.), and F32AI178919 (R.L.M.). Additional funding was provided by the BIO5 Institute Postdoctoral Fellowship from the BIO5 institute, University of Arizona awarded to R.L.M. and by the Personalized Defenses Against Disease strategic initiative from the University of Arizona awarded to J.G.P.

---

**Fig S1.**
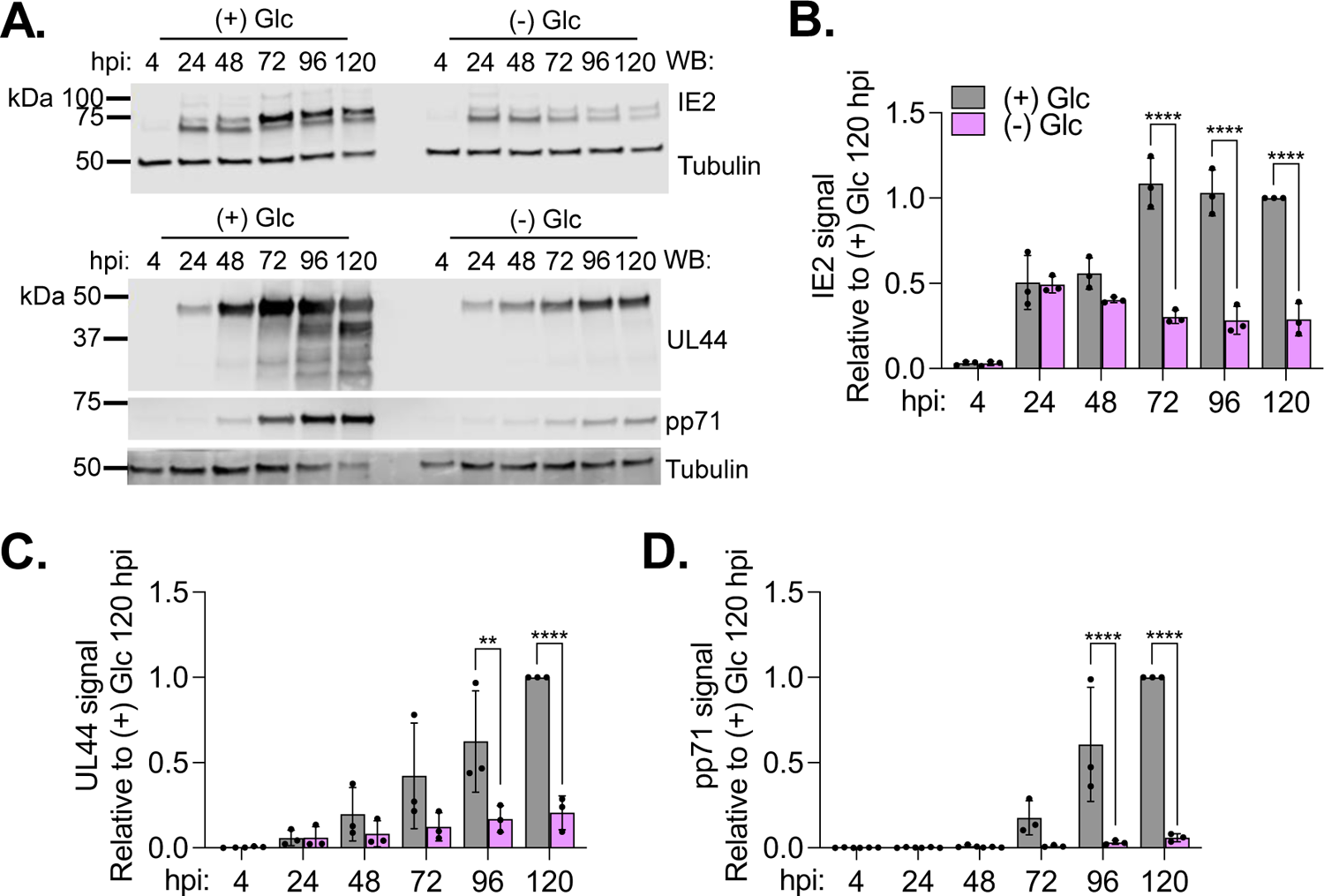
Glucose deprivation reduces early and late viral protein levels. HFFs were infected with TB40/E-GFP at a MOI 2. At 1 hpi, media was changed to glucose-replete or glucose-free DMEM. (**A**) Whole cell lysates were collected at the indicated times and analyzed by western blot. A representative blot from three biological replicates is shown. (**B-D**) Viral protein levels were normalized to tubulin levels and quantified relative to (+) Glc at 120 hpi. The graphs display three biological replicates (circles). Error bars represent SD. Two-way ANOVA with Tukey’s test was used to determine significance. *P* < 0.01, **; *P* < 0.0001, ****.

**Fig S2.**
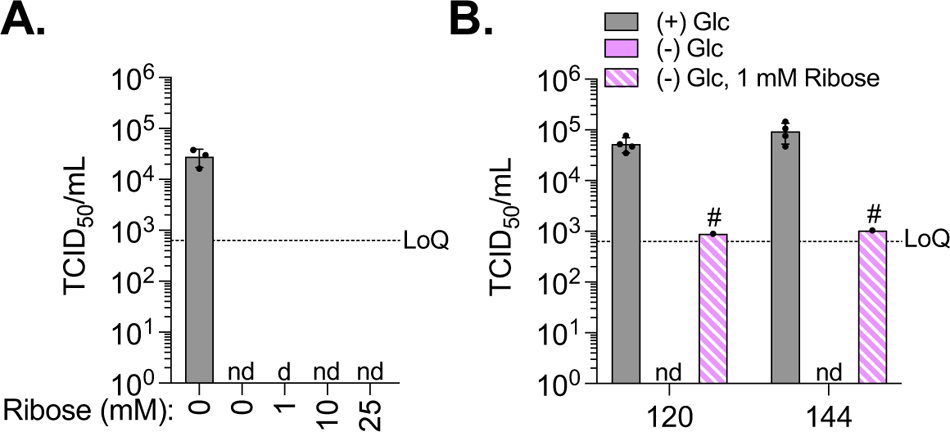
Ribose supplement restores low levels of virus production. (**A**) HFFs were infected with TB40/E-GFP at a MOI 2. At 1 hpi, media was changed to glucose-replete or glucose-free DMEM with 0-25 mM ribose supplement. Viral titers from culture supernatant at 120 hpi were measured by TCID_50_. (**B**) HFFs were infected as described above. At 1 hpi, media was changed to glucose-replete or glucose-free DMEM with 1 mM ribose from a different commercial source (Thermo Fisher Scientific). Viral titers were measured at 120 and 144 hpi by TCID_50_. LoQ = 632 TCID_50_/mL; nd = not detected; d = detected but below the LoQ. Pound sign (#) indicates that one of four biological replicates had measurable titers. One-way ANOVA with Tukey’s test was used to determine significance. *P* < 0.05, *; *P* < 0.0001, ****.

**Fig S3.**
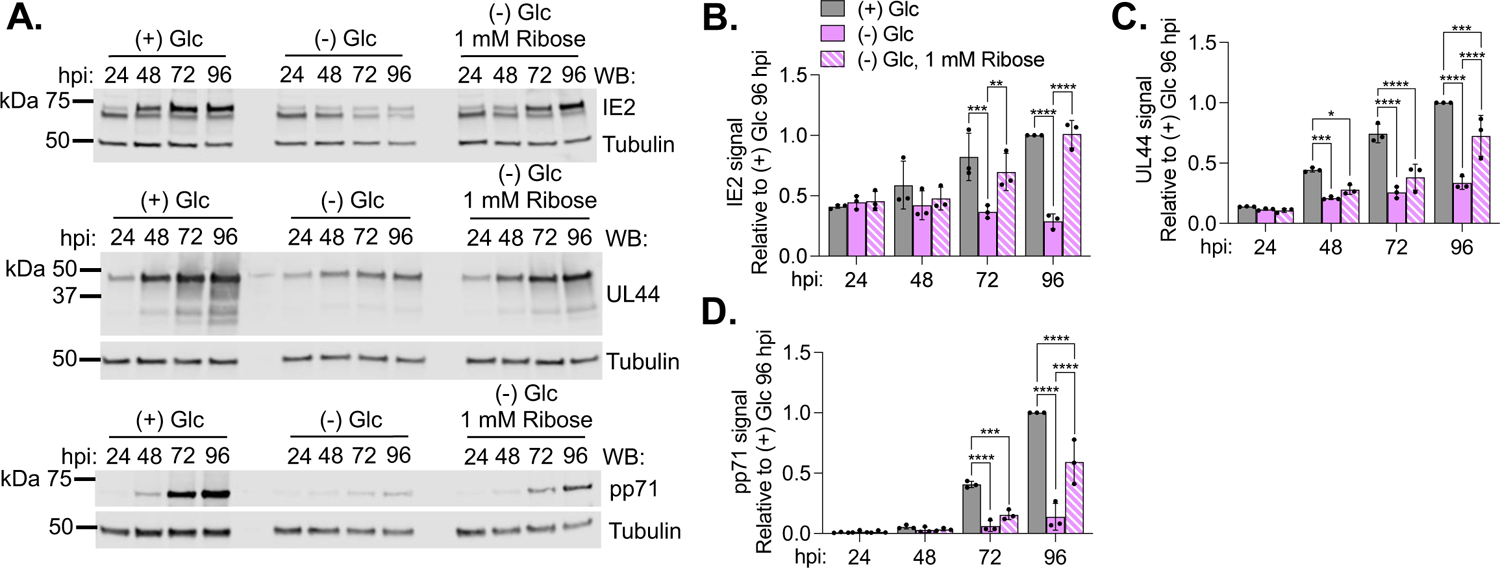
Ribose supplement partially restores viral protein levels during glucose deprivation. HFFs were infected with TB40/E-GFP at a MOI 2. At 1 hpi, media was changed to glucose-replete, glucose-free, or glucose-free DMEM supplemented with 1 mM ribose. (**A**) Whole cell lysates were collected at the indicated times and analyzed by western blot. A representative blot from three biological replicates is shown. (**B-D**) Viral protein levels were normalized to tubulin levels and quantified relative to (+) Glc at 96 hpi. The graphs display three biological replicates (circles). Error bars represent SD. Two-way ANOVA with Tukey’s test was used to determine significance. *P* < 0.05, *; *P* < 0.01, **; *P* < 0.001, ***; *P* < 0.0001, ****.

**Fig S4.**
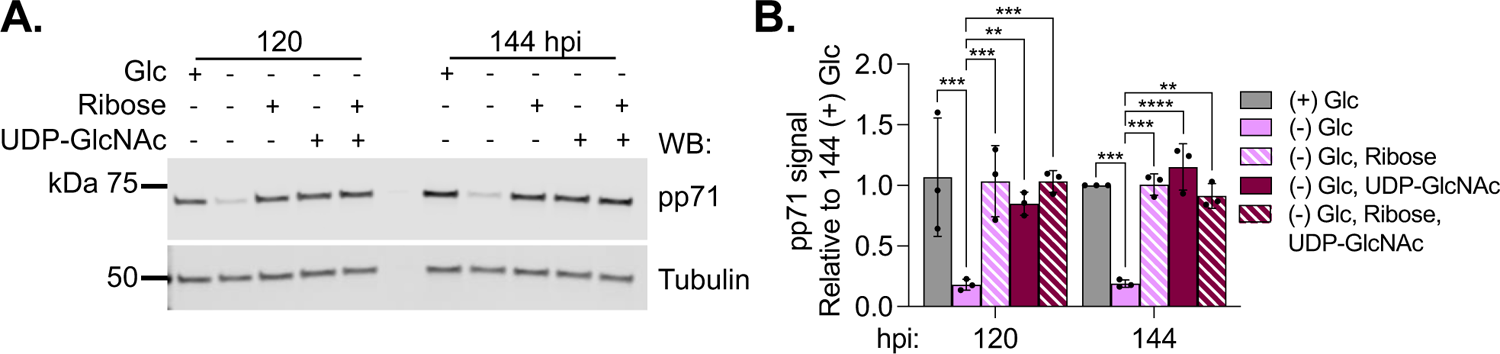
Metabolite supplement restores late protein levels. (**A-B**) HFFs were infected with TB40/E-GFP at a MOI 2. At 1 hpi, media was changed to glucose-replete, glucose-free, or glucose-free DMEM supplemented with 1 mM ribose, 100 µM UDP-GlcNAc, or dual supplement. (**A**) Whole cell lysates were collected at the indicated times and analyzed by western blot. A representative blot from three biological replicates is shown. (**B**) The levels of pp71 were normalized to tubulin levels and quantified relative to (+) Glc at 144 hpi. The graph displays three biological replicates (circles). Error bars represent SD. Two-way ANOVA with Tukey’s test was used to determine significance. *P* < 0.01, **; *P* < 0.001, ***; *P* < 0.0001, ****.

